# *VC2* regulates baseline vicine content in faba bean

**DOI:** 10.1101/2024.08.06.606773

**Authors:** Samson Ugwuanyi, Manar Makhoul, Agnieszka A. Golicz, Christian Obermeier, Rod J. Snowdon

## Abstract

Faba bean (*Vicia faba*) is a valuable legume crop desired globally for its high nutritional composition. However, the seed vicine and convicine (v-c) content reduces the nutritional quality of faba bean protein and can induce favism in individuals with glucose-6-phosphate dehydrogenase deficiency. Recently, *VC1* gene, encoding a bi-functional riboflavin protein, was reported to be responsible for initiating the biosynthetic pathway in *V. faba*. In low v-c cultivars, a 2 bp insertion in this gene results in a loss of function, but the mutation only partially eliminates v-c biosynthesis, indicating the involvement of other genes. Here, we demonstrate that a novel *V. faba* riboflavin gene, *VC2*, is responsible for the residual v-c contents in faba bean. *VC2* shares nearly identical functional domains with *VC1* and has GTP cyclohydrolase II activity, catalyzing the conversion of GTP into an intermediate molecule in the biosynthetic pathway. Gene expression analysis reveals that *VC2* contributes a minor effect to the trait, accounting for approximately 5-10% of total riboflavin gene transcripts which significantly correlates with the baseline contents in low v-c cultivars. Our results illustrate that cultivars carrying the 2 bp inactivating insertion in *VC1* still have residual v-c levels due to *VC2* activity. Furthermore, we find that *VC1* has multiple alleles and exhibits copy number variations, complicating molecular marker development. Conversely, single nucleotide polymorphisms within *VC2* provide a reliable alternative for marker-assisted selection in faba bean breeding. In conclusion, our study elucidates the complex genetic regulation of v-c biosynthesis and provides valuable insights to facilitate its elimination in faba bean.

## 1. Introduction

Faba bean (*Vicia faba* L.) is a grain legume which is globally important for its highly nutritious, protein-rich seeds. It is reported to have originated from the Mediterranean basin and subsequently spread to be cultivated across nearly all continents worldwide [1,2]. It has become increasingly popular, especially in cool-season climates where other protein crops perform poorly. The global production of faba bean stands at 5.7 million tons in 2020 [3], making it the sixth most produced pulse crop and the highest yielding legume after soybean [4]. Faba bean seeds contain up to 37% protein and are rich in micronutrients, making them a suitable source of food for humans and feed for their livestock [5,6,7]. In addition to its nutritional benefits, faba bean can improve soil fertility in association with rhizobium bacteria by fixing a significant quantity of nitrogen in the soil, which reduces the need for application of inorganic nitrogen fertilizer in subsequent seasons [4,6]. This feature is leveraged in many agricultural systems by incorporating faba bean into crop rotation or mixed cropping with others crops such as cereals [8,9]. This ecological importance of faba bean and its roles in food and feed has increased its global reputation significantly.

However, the agronomic relevance of faba bean is limited by the presence of significant quantities of vicine and convicine (v-c) in all parts of the plants [10,11]. The metabolic products of vicine and convicine, divicine and isouramil, release free radicals that cause oxidative damage to red blood cells in people with glucose-6-phosphate dehydrogenase (G6PD) deficiency, leading to acute hemolytic anemia, a condition also known as favism [11,12]. Therefore, to enhance faba bean usage and its general acceptability, reducing the v-c content to the barest minimum safe for food and feed is essential. However, the genetic basis of v-c accumulation remains to be fully elucidated, and all low v-c cultivars still carry baseline v-c levels.

Previous studies have highlighted a bimodal pattern in the v-c phenotype, primarily influenced by a major quantitative trait locus (QTL) on chromosome 1 of the faba bean genome [13,14,15]. Molecular markers flanking this locus were identified by Khazaei et al. [15], yet it was recently that the gene *VC1* was discovered within this region by Björnsdotter et al. [12]. *VC1* encodes a bi-functional riboflavin protein responsible for catalyzing the pivotal step in the v-c biosynthetic pathway. The gene has two functional domains, RibA and RibB, encoding GTP cyclohydrolase II (GCHII) and 3,4-dihydroxy-2-butanone-4-phosphate synthase (DHBPS), respectively. However, it is the GTP cyclohydrolase II domain that directly catalyzes v-c biosynthesis, via conversion of purine nucleoside triphosphate GTP into the unstable intermediates leading to v-c [12]. A 2 bp insertion in the GCHII domain leads to loss of function in low v-c cultivars. This frameshift mutation inactivates *VC1* by causing a premature stop codon, hindering the correct synthesis of GTP cyclohydrolase II. However, this mutation does not eliminate v-c completely, although it causes a significant reduction. This suggests a potential involvement of other genes or gene copies, necessitating further research to comprehensively understand the genetic factors that control v-c biosynthesis in faba bean. Recently completed *V. faba* genome assemblies and gene annotations for low and high v-c cultivars [16] provide an excellent basis to advance knowledge in this regard. The objectives of this study were to elucidate the genetic regulation of v-c content by identifying active RIBA genes and polymorphisms corresponding to changes in v-c content. Due to the residual v-c content in seeds of low v-c genotypes, we hypothesize that at least one additional locus controls v-c biosynthesis. To study this, we identified RIBA protein-coding genes by homology mapping to two faba bean reference genomes with high and low v-c content, respectively. All similar protein-encoding genes were identified and functionally analyzed across a set of well-characterized low and high v-c cultivars.

## 2. Materials and methods

### 2.1. Plant materials and cultivation

The study utilized a diverse set of well-characterized faba bean lines consisting of nine low v-c and nine high v-c genotypes with different genetic backgrounds (Supplementary Table S1). These lines represent a set of carefully selected genotypes with known characteristics and performances.

They were obtained from Norddeutsche Pflanzenzucht Hans-Georg Lembke KG (NPZ, Hohenlieth, Germany) and comprise both commercial cultivars and breeding lines beyond the fifth generation. Two seeds from each genotype were planted in 4-liter plastic pots within a pollinator-proof chamber in the greenhouse. Fresh leaves at 21 days after planting were collected from the first fully opened leaves for DNA isolation. For RNA extraction for gene expression analysis, a subset of ten genotypes was selected, including five with low and five with high v-c content. The plants were manually tripped during flowering to ensure pod-set. Fresh and immature seeds were collected from this subset for RNA isolation at two stages during seed development: early seed filling stage (ESF) (stage four) and late seed filling stage (LSF) (stage six). Specifically, seeds aged 13-16 days and 20-25 days were selected, respectively. At the ESF stage, the cleft between the cotyledons is broad, with a spherical chalazal chamber. The embryo has a butterfly-shaped appearance and is surrounded by endosperm. In contrast, at LSF, the intense green cotyledons are closely positioned and have a curved axis [17].

### 2.2. DNA extraction

Fresh leaves collected from each genotype were immediately flash-frozen in liquid nitrogen and lysed using Qiagen Tissue Lyser II (Qiagen, Düsseldorf, Germany). Genomic DNA was isolated from the leaf powder following the Doyle and Doyle [18] method.

### 2.3. RNA extraction and cDNA synthesis

Immature seeds were immediately flash-frozen in liquid nitrogen and then manually ground into powder using a mortar and pestle. RNA was isolated from 100 mg of the powdered samples using the Zymo RNA miniprep kit (Zymo Research, Freiburg, Germany) following the manufacturer’s manual. DNaseI treatment was performed to remove genomic DNA, following the procedure outlined in the Zymo RNA kit manual. RNA concentration and quality were determined using Qubit RNA assay kit (ThermoFisher Scientific, Germany) and agarose gel (1%), respectively. First-strand cDNA was synthesized using the RevertAid cDNA synthesis kit (ThermoFisher Scientific, Germany). Initially, 1 µl of Random Hexamer primer was added to 1 µg of RNA, incubated for 5 minutes at 65°C, and subsequently cooled on ice. The cDNA reaction master mix was prepared by adding 4 µl of Reaction buffer (5x), 2 µl of dNTP mix (10mM), 1 µl of Ribolock RNase inhibitor (20 U/µl), and 1 µl of RevertAid H Minus Reverse Transcriptase (200 U/µl). Reactions were carried out in a thermal cycler at the following temperature conditions: 25°C for 5 minutes, 42°C for 60 minutes, 70°C for 5 minutes, and then held at 4°C. The resulting cDNA was utilized for gene expression analysis and sequencing.

### 2.4. Homology-based gene identification

Custom databases for faba bean were established from the reference assemblies of the high v-c cultivar, Hedin, and the low v-c cultivar, Tiffany, [16] using ncbi-blast 2.12.0+. RIBA protein sequence was aligned to these databases using the tblastn function to identify all RIBA genes in both genomes. Genes exhibiting over 85% protein sequence similarity were identified, filtered and functionally analyzed in this study.

### 2.5. Phylogenetic and sequence analysis

Alignments were performed using MAFFT tool in Jalview (version 2.11.3.0). Phylogenetic analysis was conducted using the phylogeny.fr platform. The maximum likelihood method, implemented in the PhyML program (v3.1/3.0 aLRT), was employed to reconstruct the phylogenetic tree while TreeDyn (v198.3) was used for tree rendering. Amino acid sequences of RIBA proteins for chickpea (XP_004485599), lupin (KAE9591829), grass pea (CAK6822722), lotus (XP_057429064), Medicago (XP_003593237) and pea (XP_050880829) were downloaded from NCBI database. Functional domains of RIBA proteins were predicted using PsiPred workbench (http://bioinf.cs.ucl.ac.uk/psipred/).

### 2.6. Primer design and synthesis

All primers for the experiments were designed using Primer3 plus and subsequently synthesized by Microsynth AG (Balgach, Switzerland). For each primer, gene sequences from the two reference genomes were aligned to identify conserved regions with priming efficiency, as predicted by Primer3 Plus.

### 2.7. PCR Validation of *VC1* and *VC2* genes in faba bean

All *VC1* variants and *VC2* were validated in faba bean genotypes using selective PCR amplification. PCR reactions were set up in a final volume of 25 µl, including 12.5 µl of GoTaq Hot Start Green Master Mix, (Promega, Madison, WI, United States), 1.25 µl of 10 µM forward and reverse primers (Supplementary Table S2), 1.5 µl of genomic DNA and 8.5 µl of MilliQ water. The reactions were carried out in a T100 Thermal Cycler (Bio-Rad Laboratories, Hercules, CA, United States) with the following conditions: 94 °C for 2 minutes, 35 cycles of denaturation at 94 °C for 30 seconds, annealing at 60 °C for 30 seconds and extension at 72 °C for 40 to 90 seconds depending on the size of amplicon, followed by final extension for 5 minutes at 72 °C. Amplicons were separated on a 1% agarose gel and visualized under UV light.

### 2.8. *VC1* copy number determination by quantitative PCR (qPCR)

*VC1* copy number was determined by quantitative PCR in 10 µl final volume, containing 5 µl 2x SYBR Green master mix (ThermoFisher Scientific, Germany), 1 µl 10 µM forward primer and 1 µl 10 µM reverse primer (Supplementary Table S2), 1 µl of DNA and 2 µl of MilliQ water. ELF1A was used as the reference gene for normalization [19]. There were three technical replicates for each genotype, as well as triplicates of water samples serving as no-template controls. Quantitative PCR was done using StepOneplus (ThermoFisher Scientific, Germany) with the following temperature conditions: 95 °C for 10 minutes, 40 cycles of denaturation at 95 °C for 15 seconds followed by 60 °C for 1 minute. Relative quantification was determined by delta-delta CT (ΔΔCt) method [20].

### 2.9. Relative quantification of *VC1* and *VC2* expression levels by reverse-transcription quantitative PCR (RT-qPCR)

The expression levels of *VC1* and *VC2* were determined from cDNA synthesized from the previous step. The reaction mixture consisted of 5 µl of 2x SYBR Green master mix (ThermoFisher Scientific, Germany), 1 µl of 10 µM forward primer and 1 µl of 10 µM reverse primer (Supplementary Table S2), 1 µl of cDNA, and 2 µl of MilliQ water. The reference genes, CYP2 and ELF1A, were used for normalization as they were previously reported to be stably expressed in faba bean [19]. As controls, three water samples were included as no-template controls and each sample had two biological and three technical replicates. Quantitative PCR was done using StepOneplus (ThermoFisher Scientific, Germany) as described above for copy number quantification and relative transcript levels were determined by delta-delta CT (ΔΔCt) method [20].

### 2.10. *VC1* and *VC2* cDNA sequencing

The cDNA samples, synthesized as described above, were used for selective amplification of *VC1* and *VC2*. Forward and reverse primer pairs specific to *VC1* and *VC2* were used (Supplementary Table S2). The PCR conditions were as described previously in the gene validation section, except that the annealing temperature was set at 58°C. Subsequently, the resulting amplicons were sent for sequencing at Microsynth AG (Balgach, Switzerland). For each genotype, samples from the two developmental stages were sequenced.

### 2.11. KASP marker assay development and SNP genotyping

Genotyping was performed using Kompetitive Allele-Specific PCR (KASP) assay technique for polymorphisms within *VC1* gene. *VC1* genes from the two reference assemblies were aligned to identify possible single nucleotide polymorphisms. For identified variants, allele-specific primers were designed. To detect the presence of 2 bp insertion in *VC1* (SNP08) within our faba bean set, primers were designed where A1, binds to the wild-type allele, and A2, binds to the mutant allele. The common primer used was C1. Detailed information on all *KASP* markers can be found in Supplementary Table S3. The KASP marker assay procedure was conducted according to the methodology outlined in the study by Makhoul and Obermeier [21].

### 2.12. Data analysis

All experiments were repeated at least twice. The resulting qPCR data from copy number quantification and gene expressions were analyzed using Microsoft Excel and R (version 4.3.2). The library packages ggpubr and ggplot2 were used to generate plots using R studio. Statistical differences were inferred using t-test for two groups or one-way ANOVA for more than two groups.

## 3. Results

### 3.1 Multiple gene models encode RIBA proteins in the *Vicia faba* genome

In this study, we investigated the role of the different genes that encode bifunctional RIBA enzymes responsible for catalyzing the initial step in the v-c biosynthetic pathway. We employed a homology-based method to align RIBA protein to the recently assembled faba bean reference genomes, Hedin, high v-c cultivar, and Tiffany, low v-c cultivar [16]. Our analysis revealed the presence of multiple gene models encoding RIBA protein in the faba bean genome. In Hedin, we identified four such genes on chromosome 1 and contig_8341, namely Vfaba.Hedin2.R1.1g485480, Vfaba.Hedin2.R1.1g485520, Vfaba.Hedin2.R1.1g485560, and Vfaba.Hedin2.R1.Ung108560. Tiffany, on the other hand, had five genes on chromosome 1, including Vfaba.Tiffany.R1.1g399960, Vfaba.Tiffany.R1.1g400000, Vfaba.Tiffany.R1.1g400040, Vfaba.Tiffany.R1.1g400120, and Vfaba.Tiffany.R1.1g400280 (Figure 1a; Supplementary Table S4). Comparative analysis indicated that three of these genes from Hedin and four from Tiffany shared significant identity (>96%) with the *VC1* gene previously reported by Björnsdotter et al. [12] (Figure 1b). Therefore, we classified these genes as copies of *VC1*. The different *VC1* copies exhibited contrasting structural variations that grouped into three major variants, defined by their structural characteristics. *VC1A* had a tandem duplication of a 65 bp in intron 3 and *VC1B* had a partial deletion in this intron, while *VC1C* lacked the entire intron (Figure 1c). There were three copies of *VC1A* in the Hedin assembly and two copies each of *VC1B* and *VC1C* in Tiffany. As expected, one copy each of *vc1b* and *vc1c* carried a 2 bp frameshift insertion in exon 6, as reported by Björnsdotter et al. [12], which renders the gene product non-functional. This insertion was absent in *VC1A*. The modification in intron 3 is associated with a three-nucleotide difference in exon 3 and can distinguish *VC1C/vc1c* from *VC1A* and *VC1B/vc1b*. Beside these variations, *vc1b* and *VC1C* share similar alleles at most SNP positions. Additionally, there are SNPs within exon 4 and 5 that can distinguish other *VC1* variants (Supplemental Figure 1).

**Figure 1:**
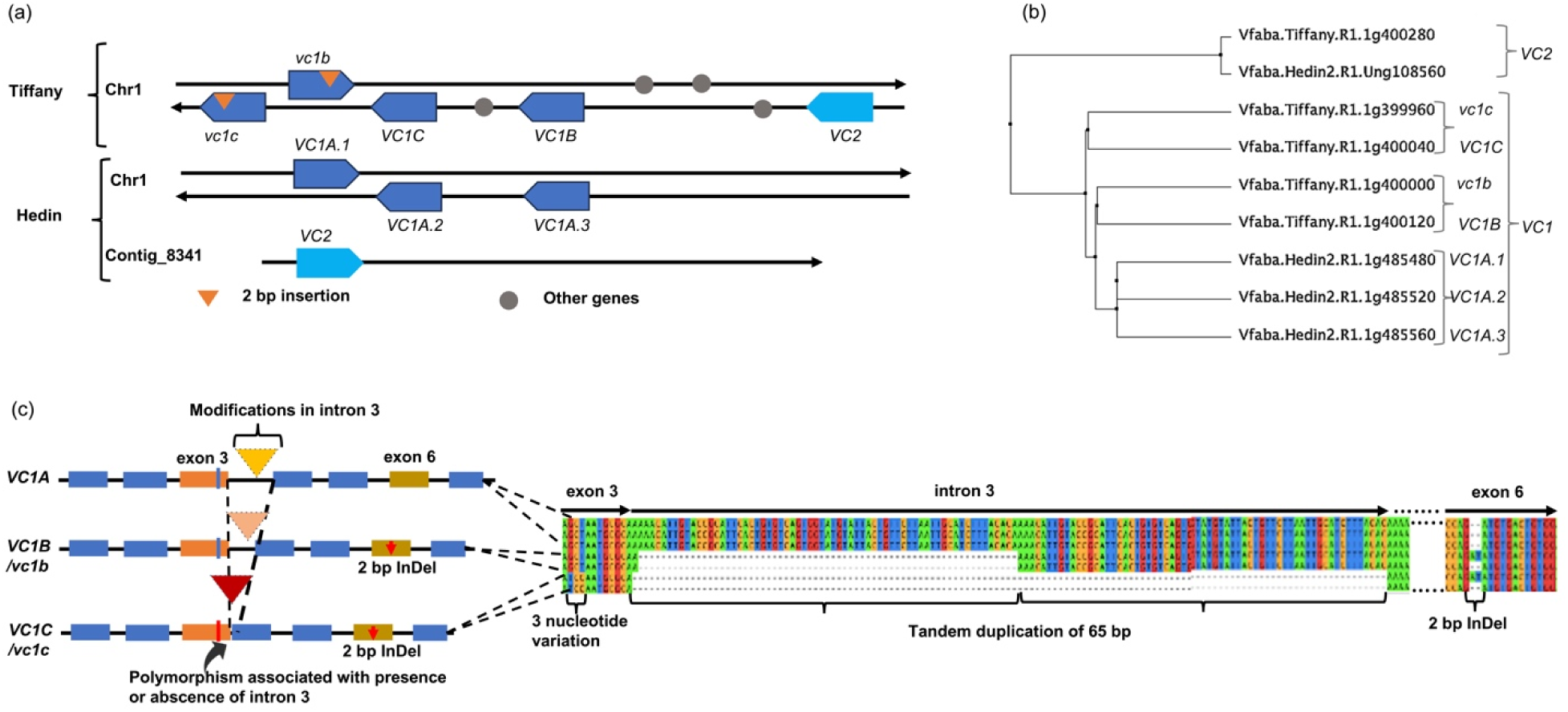
Identification and analysis of RIBA genes. (a) Configurations of RIBA genes from two reference assemblies. The direction of arrows shows the orientation of the genes on the top and bottom strand of chromosome 1 and Contig_8341. (b) Phylogenetic tree showing the genetic relationship among RIBA genes in *V. faba*. (c) Major structural variations in *VC1* genes. *VC1* variants were structurally differentiated by modifications in intron 3. These modifications are associated with a 3-nucleotide variation in exon 3.

The fourth and fifth gene models identified in Hedin and Tiffany, respectively, shared only approximately 65% sequence similarity with *VC1* and 99% sequence similarity with each other, suggesting that they represent a newly discovered *VC1*-like gene. While there were structural variations between *VC1* and this gene, these differences occurred primarily in non-coding regions, with the coding regions displaying a high degree of similarity (>87%). As a result, they shared over 90% similarity in their predicted protein sequences.

Analysis of protein sequences of this gene showed presence of two functional domains, RibA and RibB, as in other bifunctional riboflavin proteins, and highly identical to *VC1* RIBA protein domains. Subsequently, we analyzed bifunctional RIBA protein homologs from other legume crops in the Fabaceae family including chickpea, lupin, grass pea, lotus, Medicago and pea. Comparison of the amino acid sequences encoded by this newly identified *V. faba* gene, *VC1* genes and RIBA genes from these legumes revealed high similarity among these RIBA proteins and indicated the conservation of all key amino acid residues required for catalytic activities. All the proteins shared a high identity in the two catalytic domains, except *vc1b* and *vc1c* which differed significantly in the second domain due to a 2 bp insertion, altering the reading frame and causing a premature stop codon (Figure 2a-b). Hence, the high degree of similarity among these proteins, especially within their functional domains, suggests that their roles as RIBA proteins are well conserved. We therefore denominated this second RIBA gene in *V. faba* as *VC2*, since it represents a second v-c locus in *V. faba*.

**Figure 2:**
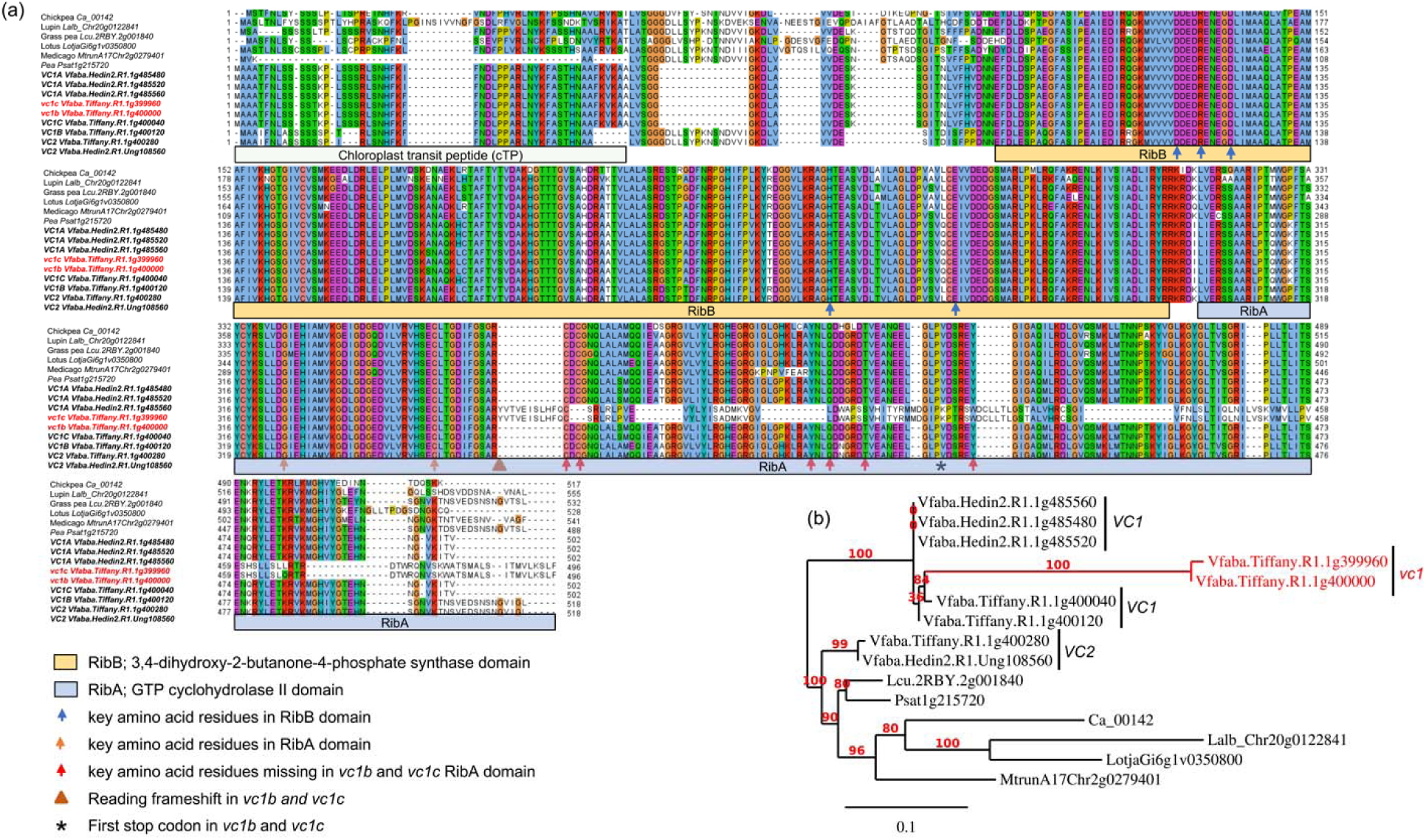
Comparison of amino acid sequences and phylogeny of RIBA proteins. (a) Alignment of amino acid sequences and (b) phylogenetic tree of RIBA proteins from *V. faba* and related legume crops in Fabaceae family, including chickpea, lupin, grass pea, lotus, Medicago and pea. Major variations lie within the domain encoding chloroplast transit pepetide (cTP). RibB and RibA domains of the proteins show high similarity, except the protein sequences encoded by *vc1b* and *vc1c* which show variations in the RibA domain due to a frameshift mutation. The functional domains are shown underneath the alignment as predicted using PsiPred workbench. Predicted key residues based on Björnsdotter et al. (2021) are indicated with different colours of arrows.

We validated the existence of all the genes in a set of 18 faba bean genotypes. First, we selectively amplified the genes using PCR. The results indicated that *VC2* was present in all faba bean genotypes. *VC1A* was present in most high v-c genotypes, while *VC1B* and *VC1C* were found in all low v-c genotypes but were also carried by some high v-c genotypes (Figure 3a). It was also observed that all three *VC1* variants could be present in a single genotype, as observed in the low v-c genotype, NPZ-FB-73, and the high v-c genotype, NP-FB-143. Secondly, the presence of mutants and wild types of *VC1B* and *VC1C* in the Tiffany genome assembly suggests that low v-c genotypes may still carry functional copies of *VC1*. To confirm the hypothesis that both mutant and wild-type copies of *VC1* are carried by low v-c genotypes, we employed a KASP assay targeting the 2 bp insertion in exon 6. KASP primers were designed with which allele 1-specific primers bind to the wild-type variant while allele 2-specific primers bind to the mutant. Subsequently, it was observed that most high v-c genotypes (7 out of 9) were homozygous for the wild-type variant, whereas all nine genotypes with low v-c and two high v-c genotypes were heterozygous, indicating the presence of both variants (Figure 3b-c). There was no homozygous call for the mutant, revealing that no faba bean genotype carried only the mutant. We extrapolated this finding to a large faba bean diversity panel comprising 97 genotypes to confirm this hypothesis. It was observed that all low v-c genotypes, as well as some high v-c genotypes, carry a functional *VC1* allele in addition to the non-functional *vc1* allele (Figure 3d).This result confirmed the hypothesis that all low v-c faba beans carry multiple *VC1* copies, including both functional and non-functional variants. Additionally, some high v-c faba beans exhibit the same characteristic.

**Figure 3:**
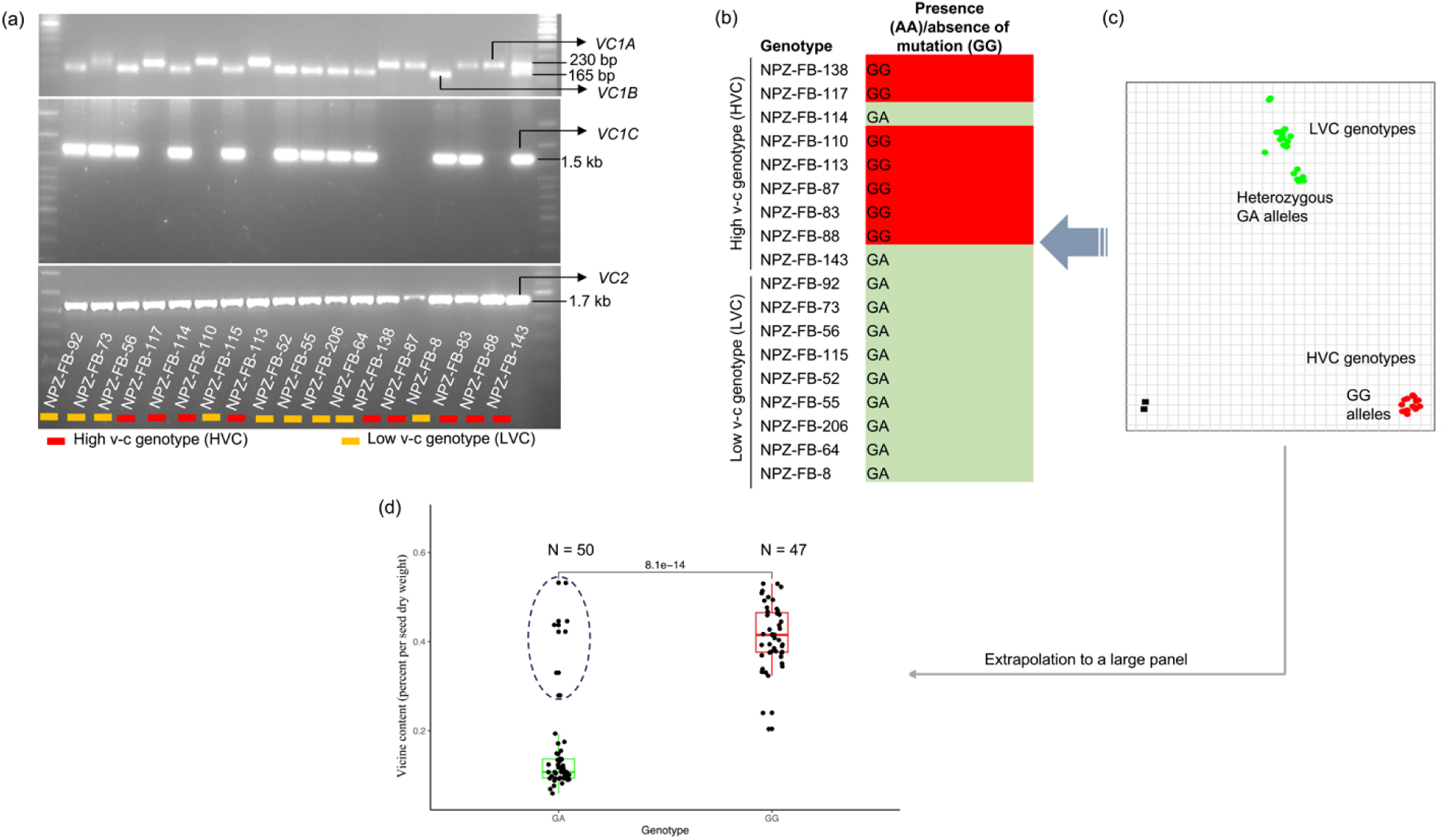
Validation of the presence of RIBA genes and the mutation in *VC1*. (a) Selective amplification by PCR showing the presence of *VC1* and *VC2* variants in a set of 18 faba bean lines. (b) Allele calls and (c) allelic discrimination plot for the 2 bp insertion in exon 6 of *VC1*. Allele GG represents the absence of insertion (wild type), while allele AA represents the presence of insertion (mutant). Heterozygous cluster (GA) indicates the presence of both the wild type and mutant variants. (d) Boxplot illustrating the v-c contents for two groups: *VC1* wildtypes (GG) and genotypes carrying both the mutant and wildtype (GA) in a panel of 97 genotypes. High v-c genotypes carrying GA alleles are enclosed within a circle.

### 3.2 *VC1* exhibits copy number variations and is dosage-insensitive

The presence of multiple copies of *VC1* in faba bean genotypes suggests a potential variable copy number for the gene. As a result, we determined *VC1* copy number through relative quantification by qPCR. The result confirmed that *VC1* shows copy number variations, with gene copies ranging from 2 to 5 across 18 faba bean genotypes (Figure 4a). Unexpectedly, low v-c genotypes tended to carry higher copy numbers than high v-c genotypes. This observation could mean that some of the gene variants are not functional or not expressed as was observed for the faba bean hilum colour locus [16]. Therefore, we assessed the transcription activities of *VC1* and *VC2* genes by quantifying their relative expression levels using RT-qPCR. A subset of ten faba bean cultivars with varying v-c contents were selected, including genotypes carrying all possible *VC1* haplotypes. Furthermore, our focus was on whole seeds at the early stage of seed development, in which *VC1* expression is known to be highest [12].

**Figure 4:**
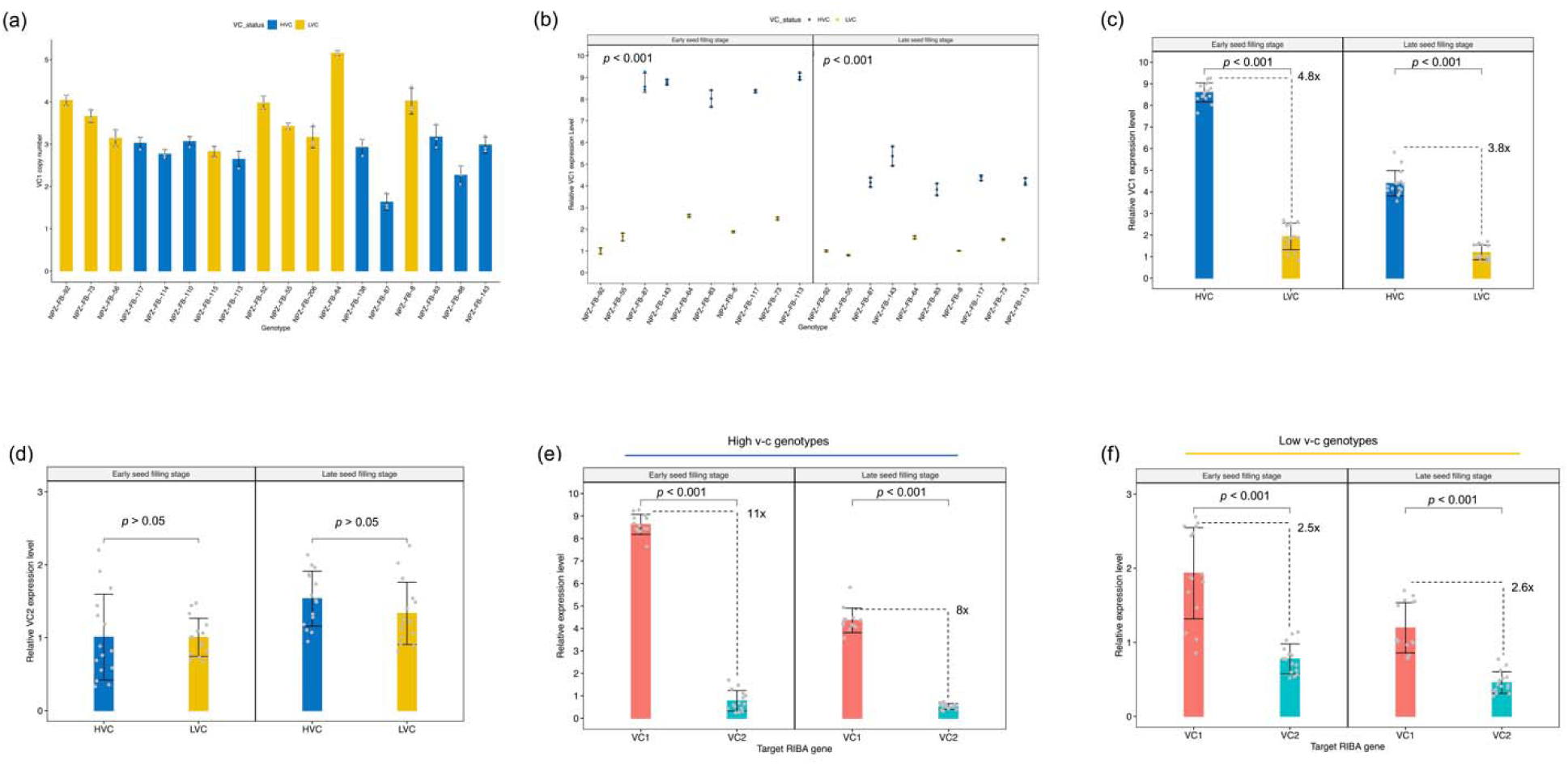
Copy number variation and transcription activities of RIBA genes. (a) *VC1* copy numbers across 18 faba bean genotypes, comprising high (HVC) and low vicine-convicine (LVC) lines (means ± SD, n = 3). (b) *VC1* relative expression levels during seed development (means ± SD, n = 3). (c) Comparisons of the relative expression levels between high and low v-c genotypes at early and late seed filling stages (means ± SD, n = 15). (d) Comparisons of relative levels of *VC2* transcripts between high and low v-c (means ± SD, n = 15). (e) Comparisons of *VC1* and *VC2* expression levels within high v-c genotypes (means ± SD, n = 15). (f) Comparison of *VC1* and *VC2* expression levels within low v-c genotypes (means ± SD, n = 15). Significance for comparisons between groups were determined using t-test.

Expression analysis revealed significantly (P < 0.001) higher *VC1* expression levels in high v-c genotypes at early seed filling and late seed filling stages (Figure 4b). High v-c genotypes showed about 5-fold and 4-fold more expression (P < 0.001) than low v-c genotypes at ESF and LSF, respectively (Figure 4c). In contrast to *VC1* differential expression, *VC2* displayed relatively consistent expression levels between high and low v-c genotype groups at both stages (Figure 4d). However, *VC1* expression was significantly higher than *VC2* expression. Within high v-c genotypes, *VC1* exhibited up to 11-fold more expression than *VC2* (Figure 4e), compared to 2.6-fold within low v-c genotypes (Figure 4f).

Importantly, despite fewer copy numbers in high v-c genotypes, the elevated expression of *VC1* suggests that it may not be sensitive to dosage. To validate this, we conducted a correlation analysis between *VC1* gene copy number and gene expression. Subsequently, we observed a negative correlation (Supplemental Figure 2a), indicating that higher copy number correlated with lower expression level. This negative correlation seems to be attributed to a shared genetic origin among low v-c genotypes, rather than interactive effects of variants resulting from antisense regulation. To verify this observation, we used a SNP genotyping dataset to trace the lineage of our low v-c lines across a broader panel of 347 faba bean genotypes. These genotypes were genotyped using the faba bean 50k Affymetrix chip. After filtering for missing data, minimum allele frequency, and heterozygosity, 13k polymorphic SNPs remained and were used for population structure analysis. The principal component analysis plot depicted a close clustering of the low v-c lines (Supplemental Figure 2b). This clustering pattern confirms a common genetic origin among low v-c genotypes, elucidating the observed negative correlation between *VC1* gene copy number and expression.

However, this observed correlation between high copy number and low expression suggests that not all *VC1* copies are expressed, potentially due to some regulatory mechanisms. To identify the functional variants, we sequenced cDNA fragments of *VC1*. As *VC1* variants can be differentiated by the mutations in exon 3 and 6, and SNPs within exon 4 and 5, we designed primers flanking this region and subsequently sequenced amplicons from four low v-c and five high v-c genotypes.

The analysis of cDNA sequences showed that three *VC1* copies were expressed across genotypes. In high v-c genotypes, sequenced cDNA corresponded to *VC1A* and *VC1C*, while *vc1b* was detected in low v-c genotypes (Figure 5a-b; Supplementary Table S5). Surprisingly, only one copy was expressed per genotype, regardless of the total number of copies present. For instance, low v-c genotype NPZ-FB-92 carried four *VC1* copies, two each of *VC1B* and *VC1C*, but only *vc1b* mutant was expressed, as in all other low v-c genotypes. Similarly, high v-c genotype NPZ-FB-143 carried multiple copies including all three variants but expressed only *VC1A*. This demonstrated that *VC1* is dosage insensitive. Additionally, we constructed a phylogenetic tree using *VC1* cDNA sequences for the sequenced subset. The genotypes clustered into two major groups (Figure 5c). The first group comprised only high v-c genotypes expressing *VC1A*, while the second group comprised two subgroups: low v-c genotypes expressing *vc1b* and high v-c genotypes expressing *VC1C*. This distinction within the second subgroup was primarily attributed to the presence or absence of a 2 bp insertion, suggesting it as a functional polymorphism within *VC1* capable of distinguishing genotypes based on v-c contents. We additionally sequenced *VC2* cDNA which overlaps with this *VC1* region, particularly within the GCHII domain. *VC2* cDNA sequences from both low v-c and high v-c genotypes did not carry any inactivating insertion (Supplementary Table S5).

**Figure 5:**
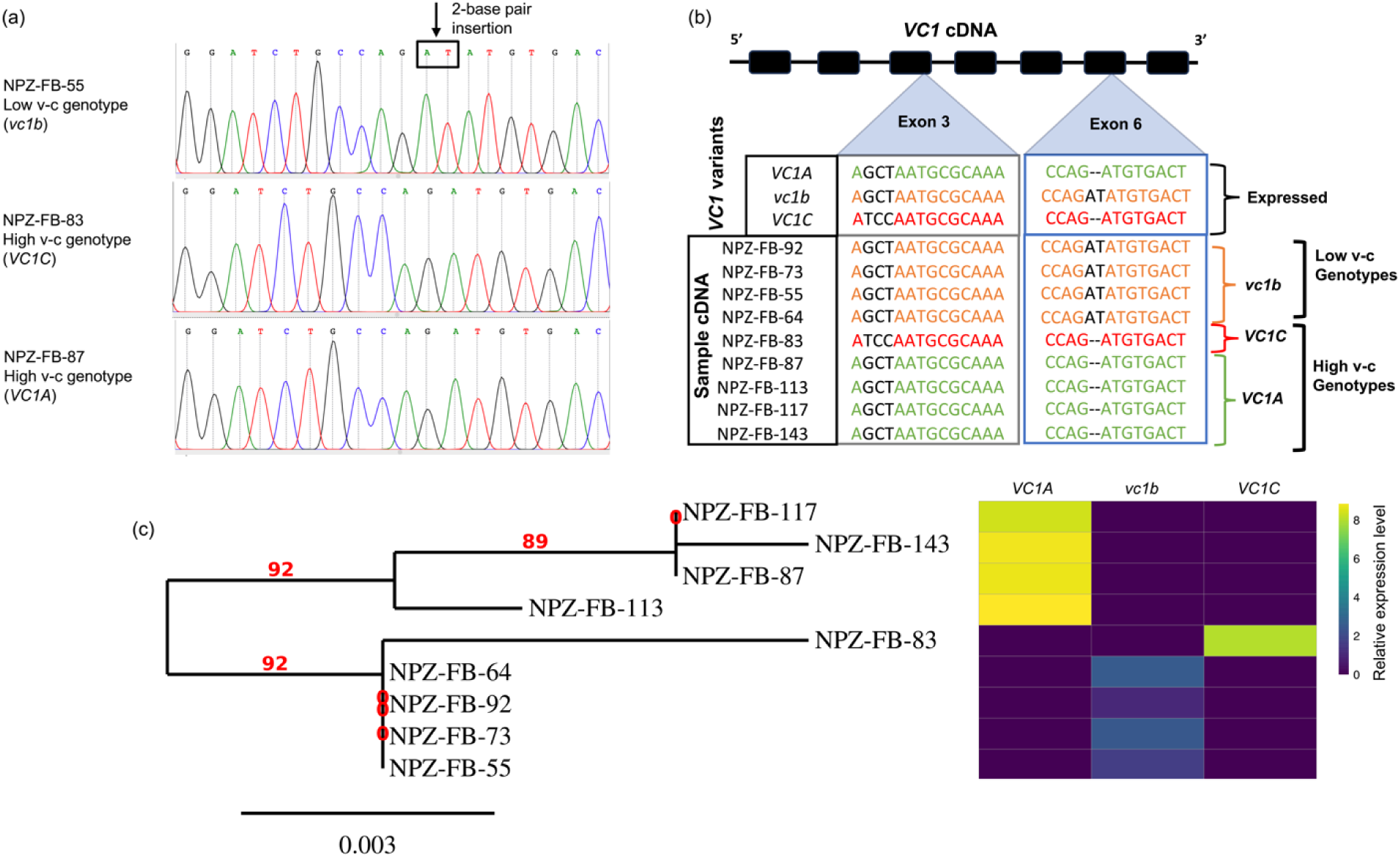
Identification of functional *VC1* variants. (a)Chromatograms obtained by Sanger sequencing of PCR fragments of VC1 cDNA show single peaks corresponding to only one copy of the gene. The position of the 2-base pair insertion is indicated by the arrow, and the nucleotides are enclosed in a box. (b) *VC1* cDNA analysis showing the expressed *VC1* variants. For each genotype, cDNA samples from two developmental stages were sequenced. In addition to the mutations shown here, single nucleotide polymorphisms in exons 4 and 5 were used to distinguish the variants (see Supplementary Table S5). (c) Phylogenetic tree showing the genetic relationships among the faba bean lines using maximum likelihood method, alongside a heatmap illustrating expression profiles for *VC1* variants.

### 3.3. *VC2* is a RIBA gene and potentially a second v-c locus in faba bean

It is evident that *VC1* is not the only locus controlling v-c biosynthesis in faba bean. Gene expression studies and cDNA sequence analysis have shown that *VC2* is a functional RIBA gene. Although there was no significant variation in expression levels of *VC2* between low and high v-c genotype groups, notable differential expressions were evident among individual faba bean lines (Figure 6a). The observed variations tended to correlate with differences in v-c contents, particularly within low v-c genotypes where *VC1* gene is inactive. Moreover, this pattern was expected since there are variations in v-c contents within low v-c genotypes, all while characterized by inactive *VC1*. Hence, this observed variation within low v-c lines might be attributed to differential *VC2* expression, as shown by high significant correlation coefficient between *VC2* expression and vicine contents (Figure 6b).

**Figure 6:**
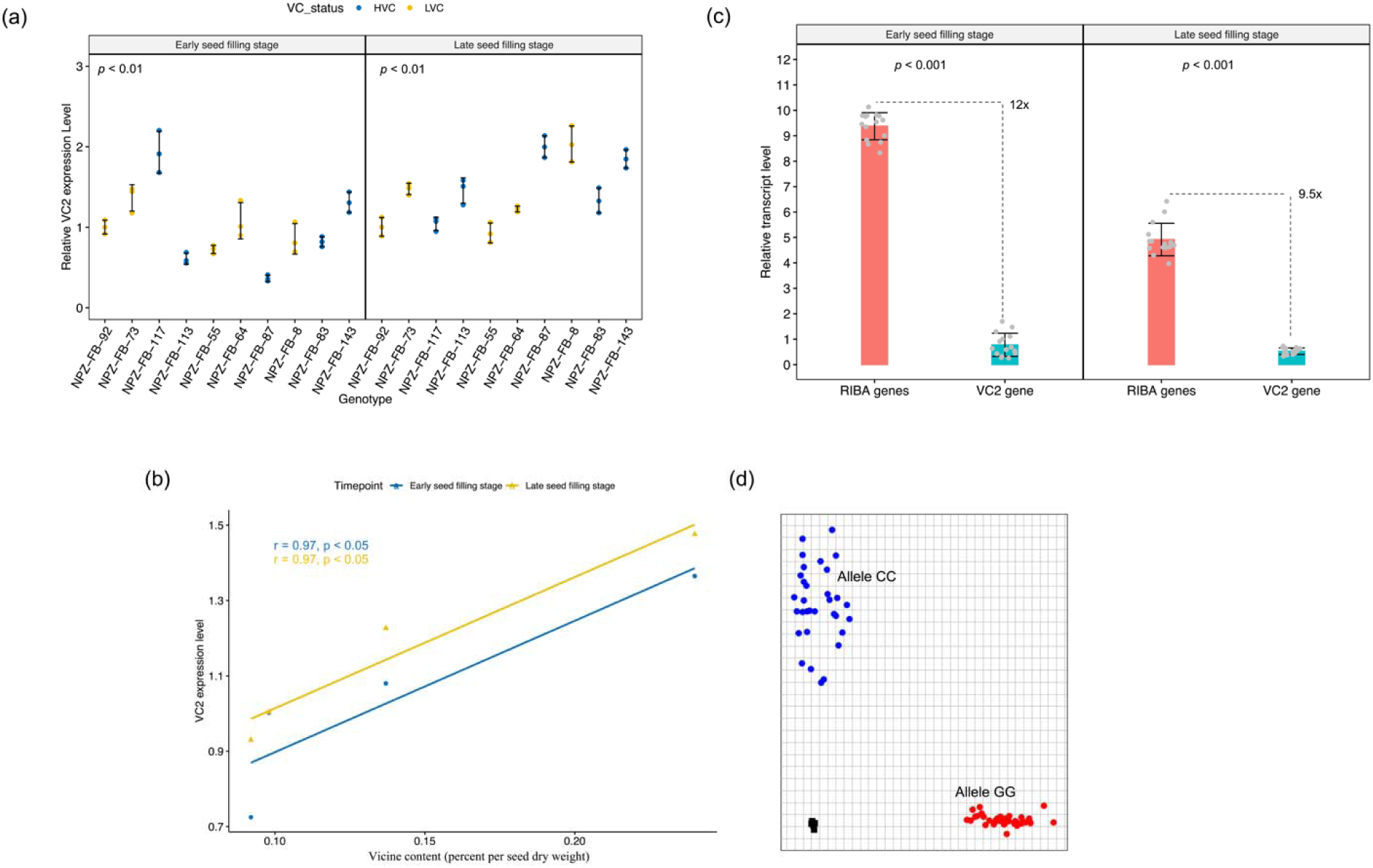
Analysis of *VC2* activity on vicine and convicine biosynthesis. (a) Relative expression levels of *VC2* across faba bean lines (means ± SD, n = 3). (b) Correlation between *VC2* expression level and vicine and convicine (v-c) contents within low v-c genotypes (means ± SD, n = 3). (c) Proportion of *VC2* expression relative to the combined RIBA gene (*VC1* and *VC2*) expression in high v-c lines at early and late seed filling stages (means ± SD, n = 15). Significance for comparisons between groups were determined using t-test. (d) Allelic discrimination plot for a SNP within exon 4 of *VC2* that segregates with the phenotype, where alleles GG and CC correlate with low and high v-c, respectively.

Furthermore, when comparing the proportion of *VC2* expression relative to the combined expression of both *VC1* and *VC2* in high v-c cultivars, we observed an average transcript level at least 9.5-fold lower, ranging up to 20-fold (Figure 6c). Assuming that only *VC2* is responsible for the observed v-c contents in low v-c genotypes, this represents about 5-10% of v-c contents in high v-c cultivars. This proportion of variation corresponds to observed phenotypic differences in v-c content between low and high v-c genotypes. In addition, earlier results from our gene expression analysis revealed that *VC2* expression levels are significantly lower than *VC1* expression. This observed expression pattern suggests that *VC2* plays a minor role in v-c regulation, and its contribution becomes evident when *vc1* is inactivated by a frameshift mutation in low v-c genotypes. Moreover, single nucleotide polymorphisms within exon 4, 5 and 6 of *VC2* segregate fully with v-c content and can differentiate low from high v-c genotypes (Figure 6d). As earlier noted, analysis of protein sequences of this new gene revealed two functional domains, RibA and RibB, similar to the functional VC1 RIBA protein domains, and conserved across legumes like chickpea, lupin, and pea. This high similarity and conservation of key amino acid residues among these proteins indicate that their roles as RIBA proteins are well conserved (see Figure 2a). Taken together, this demonstrates that *VC2* is a second functional v-c locus in faba bean.

### 3.4 Genetic variations in v-c contents involve polymorphisms associated with differential expression of *VC1* and *VC2* genes

It is commonly observed that high v-c faba beans exhibit substantial variations in v-c levels, whereas low v-c faba beans show much narrower variation, significantly below this threshold [22,23]. This suggests that a single mutation, such as the 2 bp inactivating insertion in *VC1*, may not be the sole cause behind these phenotypic differences. According to our findings, we propose that polymorphisms associated with differential expression of *VC1* and *VC2*, along with the 2 bp inactivating insertion in *VC1*, contribute to v-c content variations (Supplemental Figure 3). Our results indicate that genotype-specific differential expression of *VC1* and *VC2* leads to considerable variations in v-c contents. For instance, *VC1* expression levels among faba beans can differ up to five-fold. When combined with varying *VC2* transcript levels, this results in wider variations within high v-c genotypes. Conversely, in low v-c genotypes, the 2 bp frameshift mutation completely inactivates *VC1*. Consequently, *vc1* transcripts fail to produce functional RIBA proteins crucial in the v-c biosynthetic pathway. In these genotypes, v-c regulation is solely reliant on *VC2* activity, resulting in narrower v-c variations well below those found in wild types.

### 3.5 Implications of multiple *VC1* copies in molecular breeding for low vicine-convicine faba bean

It was observed that three different copies of *VC1* can be expressed. High v-c genotypes can express either *VC1A* or *VC1C*, while low v-c genotypes consistently express *vc1b*. Notably, *VC1C* shared close similarity with *vc1b*, having similar alleles at most SNP positions. This similarity caused genotypes expressing these copies to cluster under a major branch as observed in Figure 5c. Consequently, the existence of multiple *VC1* copies may present a significant challenge for marker-assisted selection of v-c content in faba bean breeding. Issues such as false heterozygous calls or incomplete segregation may arise due to the presence of multiple gene copies. To investigate whether the presence of multiple *VC1* gene copies can affect marker analysis, we conducted genotyping assays using well-characterized low and high v-c genotypes. These assays involved single nucleotide polymorphic markers located within *VC1* gene, including some previously developed for breeding. Since genomic DNA (gDNA) contains all gene copies, while complementary DNA (cDNA) only contains the expressed variant, we performed genotyping using KASP assays with gDNA and cDNA as templates.

Our results revealed two key findings. Firstly, the presence of multiple gene copies could lead to bias in allele calls (Figure 7a-b; Table 1). In most cases, low v-c genotypes consistently exhibited false heterozygous signals when gDNA was used as a template. A similar trend can also occur in a few high v-c genotypes. However, when cDNA was used as the template, these genotypes were correctly identified as homozygous individuals. Likewise, in the assay targeting the 2 bp insertion, all low v-c genotypes were initially called as heterozygous when gDNA was used as the template, due to the presence of multiple copies, including both wild-type and mutant alleles. However, these genotypes were correctly called as homozygous when cDNA was used as the template. Additionally, this observation further confirms that only one copy of *VC1* is expressed in each genotype analyzed.

**Figure 7:**
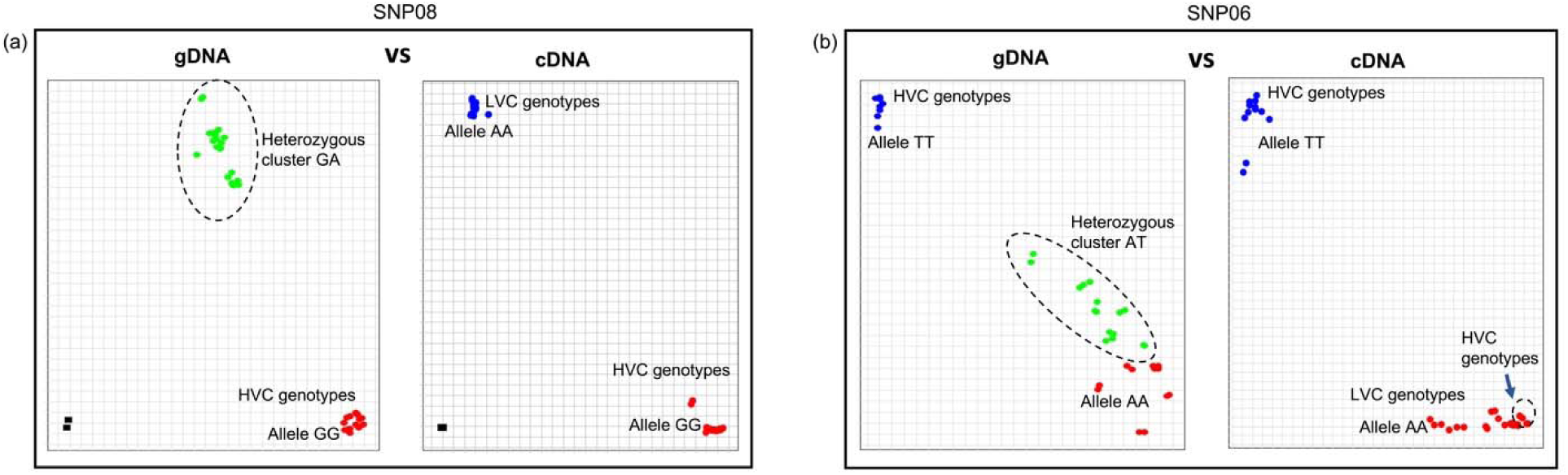
Multiple *VC1* copies can cause bias during marker analysis. Allelic discrimination plots for (a) 2 bp insertion in exon 6 and (b) an SNP within exon 5 of *VC1*. In both cases, the plots exhibited heterozygous calls for all or most low vicine-convicine (v-c) (LVC) genotypes and some high v-c (HVC) genotypes when gDNA was employed as the template. In contrast, clear homozygous calls were observed with cDNA. Additionally, some high v-c genotypes expressing *VC1C* allele can cluster with low v-c genotypes, as observed in b (see arrow).

**Table 1:**
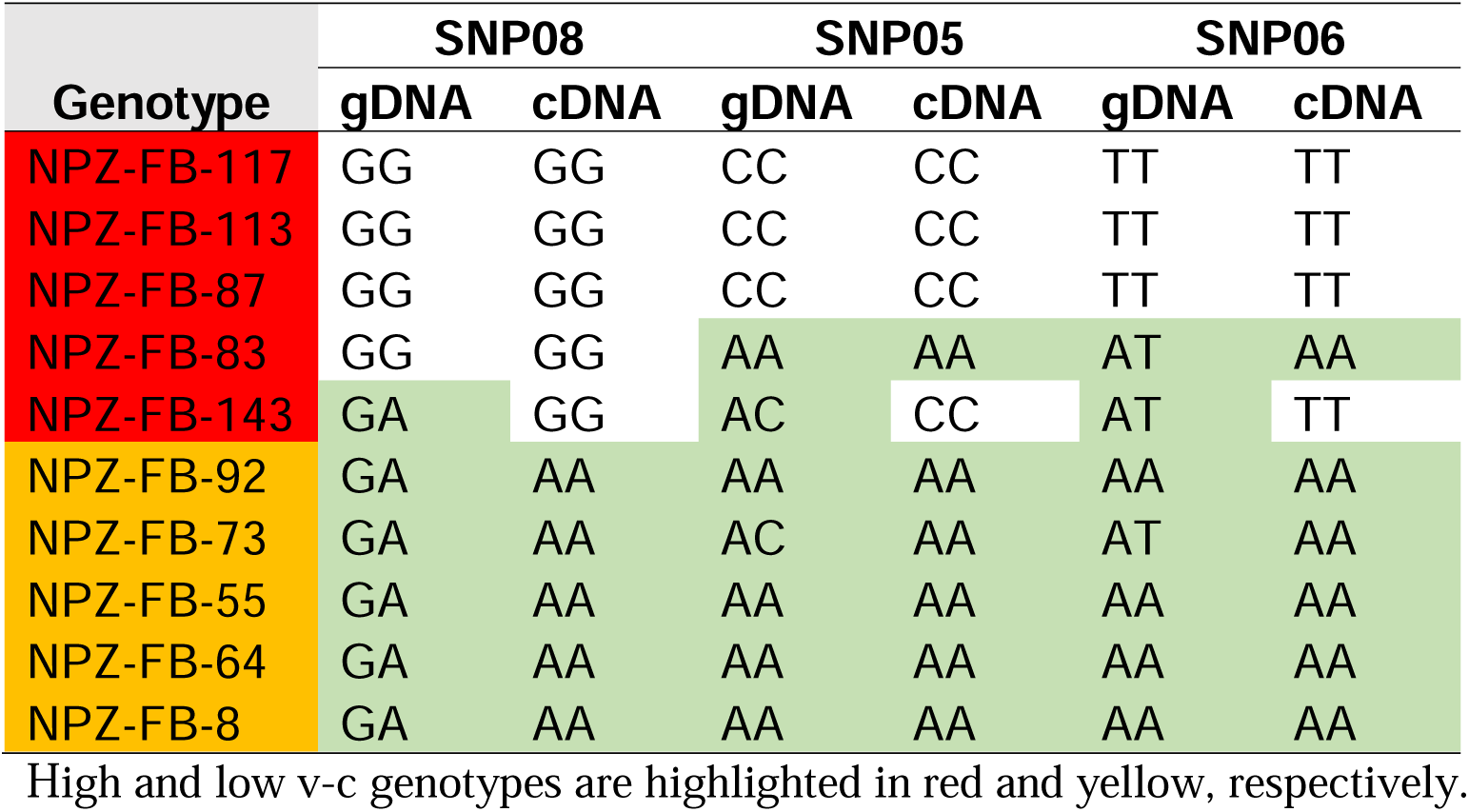
Comparison of allele calls between gDNA and cDNA for SNPs detected by KASP Assays showed that multiple copies of *VC1* often lead to inaccurate allele calls. Allele calls were also confirmed using cDNA sequences (see Supplemental Table S5).

| Genotype | SNP08 |  | SNP05 |  | SNP06 |  |
| --- | --- | --- | --- | --- | --- | --- |
|  | gDNA | cDNA | gDNA | cDNA | gDNA | cDNA |
| NPZ-FB-117 | GG | GG | CC | CC | TT | TT |
| NPZ-FB-113 | GG | GG | CC | CC | TT | TT |
| NPZ-FB-87 | GG | GG | CC | CC | TT | TT |
| NPZ-FB-83 | GG | GG | AA | AA | AT | AA |
| NPZ-FB-143 | GA | GG | AC | CC | AT | TT |
| NPZ-FB-92 | GA | AA | AA | AA | AA | AA |
| NPZ-FB-73 | GA | AA | AC | AA | AT | AA |
| NPZ-FB-55 | GA | AA | AA | AA | AA | AA |
| NPZ-FB-64 | GA | AA | AA | AA | AA | AA |
| NPZ-FB-8 | GA | AA | AA | AA | AA | AA |

High and low v-c genotypes are highlighted in red and yellow, respectively.

Secondly, we observed incomplete segregation, where some high v-c genotypes expressing *VC1C* clustered with low v-c genotypes (Figure 7b). This pattern was consistent with other *VC1*-based SNPs, except for the 2 bp insertion, which can efficiently distinguish the cultivars based on v-c contents. Beside this polymorphism, other SNPs within VC1 were ineffective in predicting v-c contents in a diverse genetic background.

## 4. Discussion

### 4.1 *VC1* is multiallelic but exhibits mono-allelic expression

Until recently, the elucidation of genetic mechanisms underlying important traits in faba bean has been hindered by the unavailability of genomic tools for this crop, which has an enormous genome in which the largest chromosome is bigger than the entire human genome. This constraint equally impeded early endeavors to identify genes responsible for v-c biosynthesis in faba bean. In the absence of reference genomes, previous investigations relied on molecular markers generated by mapping mRNA contigs from faba bean to the genomes of closely related crops, such as *Medicago truncatula*, to identify regions controlling v-c content [15]. However, the involvement of *VC1* gene in v-c biosynthesis in faba bean was established recently and a frameshift mutation in this gene results in a loss-of-function, leading to a low v-c phenotype [12].

The availability of recently assembled *V. faba* reference genomes with functional gene annotations [16] enabled us to map RIBA protein to these assemblies and identify all relevant RIBA genes associated with v-c biosynthesis. Our study revealed multiple *VC1* variants, primarily characterized by mutations in intron 3, exon 3 and 6. We found that *VC1* is affected by copy number variations, and that all low v-c genotypes carry both functional and non-functional variants, although only the non-functional variant is expressed. Previously, only two allelic forms of *VC1* were known, where low v-c faba beans carried only mutant *vc1* alleles, while high v-c faba beans carried the wild-type variant [12]. These wild-type and mutant alleles correspond to *VC1A* and *vc1b* copies, respectively. Our identification of an additional copy, *VC1C*, expressed by some high v-c lines, underscores the diversity of *VC1* genes and highlights the significance of utilizing multiple genotypes in our study. However, a major structural variation in intron 3 across the three *VC1* variants did not appear to have functional relevance as it did not associate with the expression pattern.

We observed a differential expression of VC1, consistent with previous reports between high and low v-c genotypes [12]. Surprisingly, copy number did not correlate with phenotypic expression. Although we observed a negative correlation between copy number and gene expression, this was due to the influence of low v-c genotypes sharing a single genetic source. These genotypes exhibited high copy numbers but lower expression levels. This observation was supported by cDNA sequencing which showed that each genotype expressed only one *VC1* variant despite having multiple copies in the genome. Previous reports on the transformation of hairy roots of Hedin with an additional copy of functional *VC1* did not result in increased vicine accumulation [12]. This confirms that despite copy number variation, *VC1* is not sensitive to dosage.

It is not clear what mechanisms are involved in *VC1* dosage compensation; however, certain mechanisms have been proposed for dosage compensation in genes with multiple copies which could involve complex processes that span different stages at transcription [24,25]. MicroRNAs (miRNAs) are one of the mechanisms proposed as potential mediators of dosage compensation, where they play a role in fine-tuning gene expression [25,26]. Although some studies reported that miRNAs can activate transcription and indicated a possible preference for miRNA regulation in duplicated or copy number variable (CNV) genes [27,28], Woodwark and Bateman [25] and Chang and Liao [29] presented contradictory findings, suggesting fewer CNV genes regulated by miRNAs than initially expected. Additionally, dosage compensation mechanisms, such as imprinted or monoallelically expressed genes, are widely reported. While not fully understood, imprinted genes typically deactivate one parental allele through antisense non-coding RNAs, whereas monoallelic genes express only one allele, regardless of parental origin [30]. DNA methylation is another potential mechanism widely reported in the regulation of duplicated genes. It is frequently associated with a transcriptionally inactive state [31,32], and studies have shown that modifications of DNA methylation occurring upstream of genes can impede the initiation of transcription in duplicated genes [29]. Methylation is more likely to play a role in the regulation of multiple copies of *VC1*, where methylation of the additional gene copies may impede their transcription, resulting in the expression of only one gene copy.

The regulation of *VC1* differential expression between low and high v-c genotypes and the impact of the 2 bp insertion carried by low v-c genotypes on expression levels remain unclear. Differences in the expression of *VC1* genes can also stem from variations within the regulatory elements. Elements like enhancers and silencers have the potential to amplify or suppress the gene expression of the target gene [33,34]. Upstream and downstream of *VC1* are various variants, such as short tandem repeats, InDels and SNPs. Structural differences within this region could potentially affect one or more regulatory elements. The interaction among these diverse regulatory components, their interplay with target promoters, and the involvement of epigenetic modifications intricately regulate gene expression [33,34,35,36,37].

### 4.2 *VC2* is a functional RIBA locus potentially catalyzing the biosynthesis of baseline vicine-convicine in low v-c faba bean

The variation in *VC1* expression alone does not correlate with observed phenotypic differences among faba bean genotypes. Subsequent analysis of *VC1* cDNA in this study aligns with findings reported by Björnsdotter et al. [12], that all low v-c cultivars express *vc1b* alleles carrying 2 bp insertion in exon 6 which results in a non-functional protein. This implies that *VC1* is not active in low v-c genotypes. However, as previously mentioned, this mutation only causes a significant reduction in v-c content but does not entirely eliminate it. For instance, the seed v-c content of the first low v-c genotype, initially reported by Duc et al. [38] is approximately 0.04%, which is about 1/10 to 1/20 of high v-c contents. This substantial reduction in v-c content due to a single gene mutation underscores the major role of *VC1* in v-c phenotype regulation and equally suggests the involvement of another gene with a minor effect.

In addition to the already known *VC1*, we identified an additional gene, *VC2*, encoding putative bifunctional riboflavin proteins in faba bean genome. The distinct structure of the *VC2* gene compared to *VC1* suggests that it represents a second RIBA locus in faba bean. *VC2* is a bifunctional riboflavin gene with two catalytic domains, RibB and RibA, encoding DHBPS and GCHII enzymes, respectively. RIBA enzymes are known for their involvement in the riboflavin biosynthetic pathway in plants [39]. However, Björnsdotter et al. [12] demonstrated that RIBA enzymes are also involved in v-c production where v-c are synthesized in a three-step pathway that starts from the GTP cyclohydrolase II function of RIBA proteins. While *VC1* and *VC2* share nearly identical functional domains, *VC2* lacks the inactivating insertion present in *vc1b* mutant. Detailed analysis of *VC1* and *VC2* amino acid sequences showed conservation in all key amino acid residues necessary for catalytic activities, including those required for binding zinc ions essential for GCHII activity as demonstrated by Kaiser et al. [40]. These key amino acid residues are well conserved in *VC1* and *VC2* and other RIBA proteins within Fabaceae family but lacking in *vc1b*.

Subsequent functional analysis demonstrates that *VC2* is a functional RIBA gene and represents a second v-c locus in faba bean. This hypothesis was supported by our expression analysis, which revealed lower *VC2* expression levels relative to combined RIBA gene transcripts in high v-c genotypes. This proportion of variation aligns with the observed phenotypic differences between low and high v-c genotypes, assuming *VC2* transcript accounts for the residual contents in low v-c lines.

Previous studies consistently emphasize the *VC1* locus as a major determinant in v-c regulation [2,13,14,15]. Our study substantiates this hypothesis, indicating several-fold higher expression of *VC1* relative to *VC2*. While the significance of *VC1* is well-established, the presence of *VC2* explains why mutation in *VC1* does not eliminate v-c completely in affected genotypes. This minor effect, coupled with its non-differential expression between high and low v-c lines, could explain why it remained unidentified in previous studies using transcript-based annotation [2,12]. Previous studies often targeted genes exhibiting differential expression between high and low v-c lines. In addition, transcript-based annotation methods may not efficiently differentiate *VC1* from *VC2* as coding regions are highly similar.

Our findings demonstrate that *VC2* is a functional RIBA locus responsible for the residual v-c contents in low v-c genotypes. However, it is essential to validate the involvement of *VC2* in the biosynthesis of v-c through mutant analysis. Unfortunately, the absence of reliable transformation methods for faba bean poses a significant limitation to validate gene function by knockouts.

Unlike other crops, faba bean lacks mutant collections and well-established protocols for gene knockout [41], and considerable difficulties have been encountered in developing and regenerating transgenic/CRISPR plants in this species.

### 4.3 Multiallelic nature of *VC1* makes the application of localized SNPs inefficient for marker-assisted selection in vicine-convicine breeding

Various molecular markers have been developed to facilitate breeding of low v-c faba beans [2,42]. However, most existing v-c markers target polymorphisms within *VC1* genes. While these markers have proven valuable in some contexts, their efficiencies may be limited due to multi-allelic nature of *VC1*. Our research revealed that multiple VC1 copies can lead to inaccurate allele calls, resulting in false clusters during marker analysis. This often happens as a result of the existence of closely homeologous sequences in the genome [43]. Consequently, conventional KASP assays may require optimization to improve accuracy [43]. Nevertheless, the applicability of these SNPs might be confined to specific genetic backgrounds. In a diverse genetic background, comprising all expressed *VC1* alleles, SNPs often do not segregate fully with the phenotype. Hence, these polymorphisms may not be efficient for MAS of v-c in faba bean breeding. Therefore, we strongly advise caution when utilizing *VC1*-based molecular markers for v-c content selection in breeding programs. Overreliance on only these markers for selection may inadvertently lead to incorrect prediction of v-c contents.

However, our study highlights the functional significance of the 2 bp insertion in exon 6 as a reliable polymorphism in *VC1* that effectively distinguishes genotypes based on the v-c phenotype. Optimizing the KASP assay targeting this polymorphism could significantly enhance its efficiency and specificity, particularly given the potential for inaccurate clustering due to multiple gene copies. Moreover, SNPs within *VC2* show consistent segregation with v-c contents. These SNPs can accurately predict v-c contents without any bias and are therefore recommended for breeding. This gene holds the potential to provide reliable molecular tools for marker-assisted selection of v-c content in faba bean breeding programs.

## 5. Conclusion

We have demonstrated that *VC1* has multiple copies and shows CNV. However, copy number does not correlate with gene expression and suggests a tight regulation of the gene. It was observed that multiple *VC1* variants were expressed among low and high v-c genotypes which complicates molecular marker development for breeding. Importantly, we identified a second RIBA gene, *VC2*, which shared nearly identical RIBA domains with *VC1*, and does not carry any inactivating mutation. *VC2* is functional in all genotypes and represents a second gene underlying v-c in faba bean, explaining why mutation in *VC1* does not result in zero v-c. Although the role of *VC2* gene in v-c biosynthesis must still be validated for example using editing knockouts, our results already provide significant information to facilitate elimination of v-c in faba bean.

## Acknowledgments

The authors gratefully acknowledge Stavros Tzigos and Annette Plank for their valuable technical assistance with nucleic acid extraction and plant management in the greenhouse, respectively. We thank Erick Owuor Mikwa for his insightful comments and suggestions. We also express our gratitude to Norddeutsche Pflanzenzucht Hans-Georg Lembke KG (NPZ, Hohenlieth, Germany) for providing the faba bean seeds used in the study.

## Author contribution statement

RJS and SU conceived the study; SU conducted the experiments, analyzed the data, interpreted the results, and drafted the manuscript; SU and MM performed KASP assay; CO and AAG assisted in data analysis; AAG provided gene and protein sequences. All authors revised the manuscript and approved the final manuscript.

## Data availability

The authors declare that all data reported in the paper are contained within the paper and supplementary information.

## Funding

The work was supported by DFG grant 469336000 to RJS from the German Research Society (DFG) for the International Research Training Group IRTG 2843 Accelerating Crop Genetic Gain. DFG grant 497667402 to AAG.

## Competing Interests

The authors have no relevant financial or non-financial interests to disclose.

## List of Abbreviations

cDNA: Complementary DNA
CNV: Copy number variation
DHBPS: 3,4-Dihydroxy-2-Butanone-4-Phosphate Synthase
DNA: Deoxyribonucleic Acid
ESF: Early Seed Filling
G6PD: Glucose-6-Phosphate Dehydrogenase
GCHII: Guanosine Triphosphate Cyclohydrolase II
gDNA: Genomic DNA
GTP: Guanosine Triphosphate
HVC: High Vicine-Convicine
KASP: Kompetitive Allele-Specific PCR
LSF: Late Seed Filling
LVC: Low Vicine-Convicine
miRNA: MicroRNA
NCBI: National Center for Biotechnology Information
PCR: Polymerase Chain Reaction
qPCR: Quantitative Polymerase Chain Reaction
QTL: Quantitative Trait Locus
RNA: Ribonucleic Acid
RT-qPCR: Reverse Transcription Quantitative Polymerase Chain Reaction
SNP: Single Nucleotide Polymorphism
v-c: Vicine and Convicine

## Notes

### Competing Interest Statement

The authors have declared no competing interest.

