## Supplemental Figures for "*VC2* regulates baseline vicine content in faba bean"

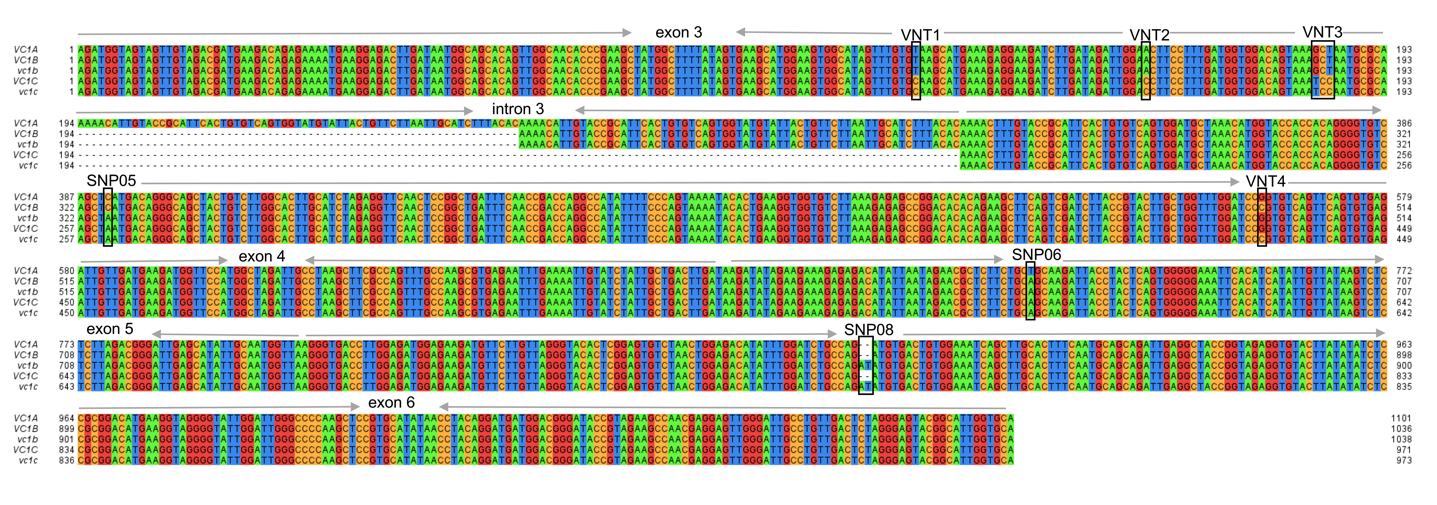
**Supplemental Figure 1:** Structural variations in VC1 genes. Alignment of reference VC1 genes from Hedin and Tiffany showing polymorphisms in the coding regions of the genes that can distinguish different variants.

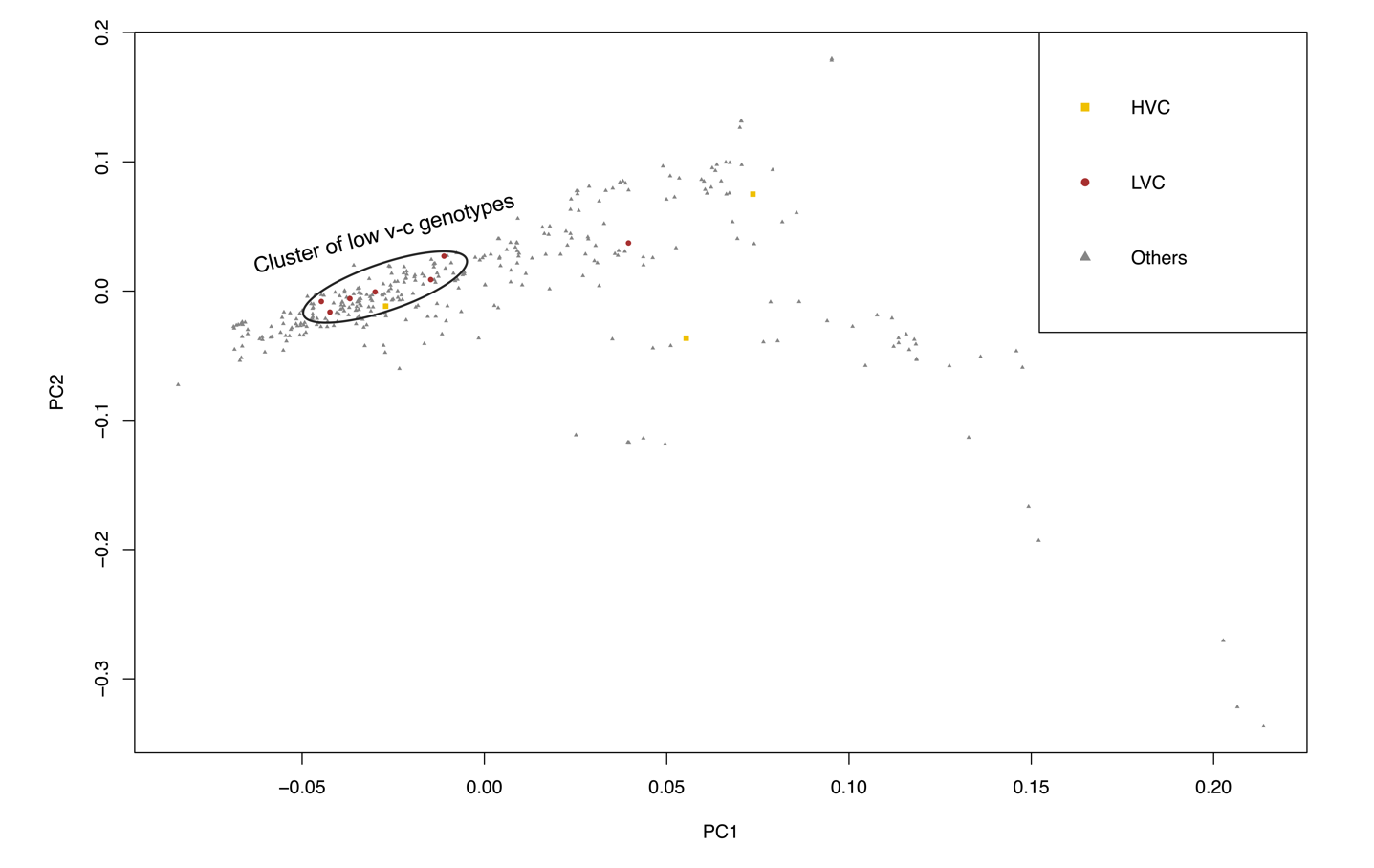
**Supplemental Figure 2**: **Tracing the source of low vicine and convicine (v-c) trait**. (a) Correlation analysis showing the relationship between VC1 copy number and VC1 expression (means ± SD, n = 3). (b) Principal component analysis showed that most low v-c genotypes cluster together as they share a single genetic origin, as shown in dark circle.

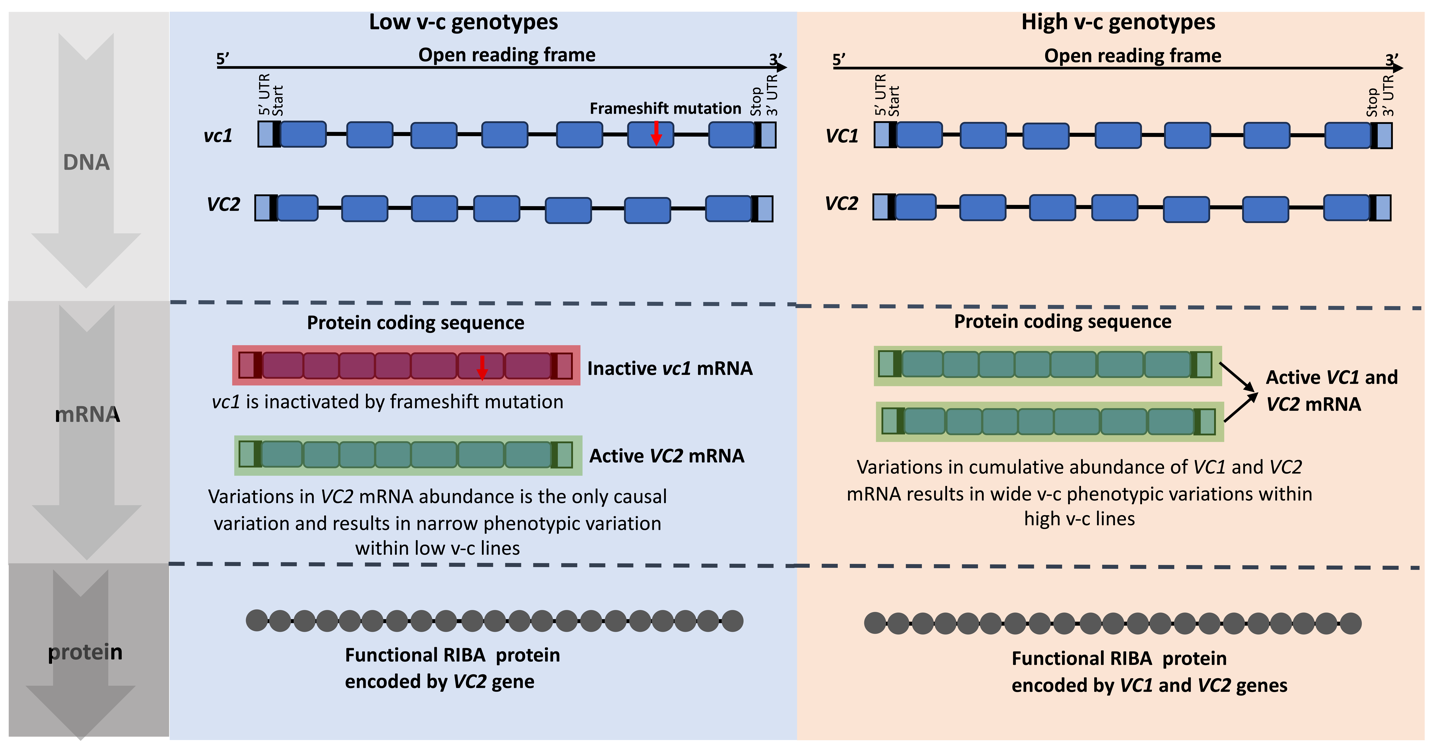
**Supplemental Figure 3**: Schematic representation of genetic variations in vicine and convicine (v-c) contents involving *VC1* and *VC2* genes.
